# Multi-Omics Study of Ancestry in Adults with Intracranial Cancers – Glioma (MOSAIC)

**DOI:** 10.64898/2026.06.24.733669

**Authors:** Melissa L. Bondy, Humaira Noor, Spiridon Tsavachidis, Kazutaka Fukumura, Quinn T. Ostrom, Kyle M. Walsh, Bo Peng, Donna Marie Muzny, Viktoriya Korchina, Burt Nabors, Lisa Norberg, Annick Desjardins, Jennifer Ritchie, Craig Horbinski, Alejandro Perez, Vasudev Tadimeti, Jacob Mandel, Margaret Wrensch, Tejus A. Bale, Irene Orlow, Jianhong Hu, Harsha Doddapaneni, Xiuping Liu, Zeineen Momin, Pryanka Motewar, Georgina Armstrong, Meghan Woods, Jonine L. Bernstein, Christopher I. Amos, Jason T. Huse

## Abstract

**Background:** Most genomic studies of adult-type diffuse gliomas have focused on predominantly European ancestry populations, limiting the generalizability of molecular classifications and precision medicine approaches. We assembled a multi-institutional glioma cohort of diverse patients to investigate how germline ancestry, molecular subtypes, and mutational processes shape tumor biology and clinical outcomes.

**Methods:** We analyzed 1,102 adults with WHO 2021–classified diffuse gliomas (IDH-mutant, 1p/19q-codeleted oligodendroglioma; IDH-mutant astrocytoma; IDH-wildtype glioma) from seven U.S. institutions. Whole-exome sequencing (WES) of FFPE tumors identified somatic alterations and COSMIC SBS v3.2 mutational signatures. Genetic ancestry was estimated from WES using 1000 Genomes reference populations. Overall survival was assessed using Kaplan–Meier and multivariable models.

**Results:** The cohort included 66.9% European (EUR), 21.1% Admixed American/Hispanic (AMR), 10.3% Admixed African (AFR), and 1.6% Asian (AS) ancestry. Survival followed expected molecular hierarchy (median overall survival: oligodendroglioma 15.7 years, astrocytoma 10.6 years, IDH-wildtype glioma 1.9 years). Within oligodendroglioma, AMR patients showed improved survival versus EUR (HR 0.67, 95% CI 0.48–0.94; p=0.011), with similar trends across subtypes. Somatic profiling confirmed canonical subtype-defining alterations and revealed higher ATRX alterations in AFR and AMR IDH-wildtype tumors compared with EUR. ATRX alterations were associated with improved survival only in AFR (p=0.003). Mutational signature analysis identified subtype-specific signatures, including therapy-associated signatures. Chemotherapy-related signatures were more frequent in EUR and AMR than in AFR.

**Conclusions:** This ancestrally diverse glioma cohort confirms established molecular classifications and identifies ancestry-associated differences in survival, somatic alterations, and mutational processes, indicating the critical need for broad representation to inform precision neuro-oncology.

**Key Points:**

- A multi-institutional glioma cohort validates subtype and survival patterns across ancestries.
- Therapy-associated mutational signatures differ by ancestry, suggesting distinct treatment-related mutational processes.
- Admixed American patients show improved survival, particularly in oligodendroglioma.

**Importance of the Study:** Most genomic studies of adult-type diffuse glioma have focused on populations of predominantly European ancestry which limits the ability to examine variation in tumor biology and clinical outcomes across populations. In this study, we assembled one of the largest ancestrally diverse cohorts of molecularly characterized adult diffuse gliomas, integrating germline ancestry inference with tumor whole-exome sequencing and mutational signature analysis. We confirm that established molecular classifications and survival hierarchies remain robust across ancestry groups. However, we also identified ancestry-associated differences in survival within specific tumor subtypes, higher *ATRX* alteration frequencies in African American and admixed American patients with IDH-wildtype tumors, and variation in therapy-associated mutational signatures across ancestry groups. These findings highlight the importance of incorporating population differences into genomic studies of glioma and provide a resource for future multi-ancestry investigations of glioma risk, tumor evolution, and treatment response, ultimately supporting more inclusive precision neuro-oncology.

## INTRODUCTION

Gliomas are the most common primary malignant brain tumors in adults and account for most central nervous system (CNS) cancers diagnosed each year in the United States. In 2026, an estimated 24,750 new cases of primary malignant brain tumors are expected, of which roughly 80% will be gliomas [1, 2]. In the latest report, annual glioma incidence is highest among non-Hispanic White individuals at 7.3 per 100,000, followed by Hispanic individuals at 5.3 per 100,000, while Asian and Black populations show similar rates, ranging from 4.3 to 4.5 per 100,000. Overall, more than 85% of gliomas in the US occur in White individuals, compared with approximately 6% in Black or African Americans and 7% in Hispanic populations [2–4]. Glioblastoma (GBM) incidence is also higher in males than females, with men experiencing a 50–60% greater risk; however, the basis of this sex difference remains poorly understood. Although access to specialty care and social determinants of health likely influence diagnosis and outcomes, the acute neurologic presentation of gliomas often leads to prompt imaging, which may reduce but not eliminate systematic underdiagnosis, particularly in populations with reduced access to care [5]. Despite variation in histology and clinical grade, the fundamental challenge in glioma management lies in the inherently invasive biology of these tumors and their resistance to available surgical and medical therapies. GBM, the most aggressive subtype, is characterized by especially rapid progression and poor prognosis [2].

Lower-grade astrocytomas, IDH-mutant (Astro) and oligodendrogliomas, IDH-mutant and 1p/19q co-deleted (Oligo), often follow more indolent clinical courses, with median survivals on the order of 7-15 years depending on subtype and treatment. In contrast, median overall survival for GBM remains approximately 12–14 months even with maximal resection and aggressive chemo-radiotherapy, and fewer than 7% of patients with GBM survive beyond five years [2, 6]. The average years of potential life lost in glioma, estimated at about 20 years per patient, exceeds that of any other major adult cancer [7, 8]. Several studies report that African American and Hispanic patients with GBM have equal or better survival than non-Hispanic White patients, even after adjustment for age, treatment, and comorbidities [9–14]. These statistics underscore the need for deeper insight into glioma pathogenesis and sources of heterogeneity to enable precision risk stratification.

Large-scale molecular profiling efforts in adult diffuse gliomas, including The Cancer Genome Atlas (TCGA), identified recurrent somatic alterations that define biologically and clinically distinct entities. Central among these are mutations in *IDH1/2* and their characteristic patterns of co-occurring alterations, such as *TP53* and *ATRX* mutations or whole-arm *1p/19q* co-deletion, which underpin the 2021 WHO classification of adult diffuse gliomas [15, 16]. Incorporation of these molecular features into the WHO CNS classification has led to a taxonomy that emphasizes three major categories of adult-type diffuse glioma: with IDH-mutant astrocytoma, which typically harbors inactivating *TP53* and *ATRX* mutations; IDH-mutant and 1p/19q-codeleted oligodendroglioma, which is also essentially defined by *TERT* promoter mutations; and IDH-wt gliomas, which in most cases show aggressive GBM-like behavior despite considerable molecular heterogeneity [1].

The majority of molecular studies that have shaped this understanding of glioma biology are drawn predominantly from cohorts of European or East Asian ancestry. As a result, patients from other racial ethnic groups, including African American and Hispanic populations, remain substantially underrepresented in genomic, epidemiologic, and clinical glioma research, and the molecular characteristics of gliomas in these groups are still poorly defined [17, 18]. This lack of diversity has constrained discovery and limited the generalizability of biological insights, diagnostic biomarkers, therapeutic targets, prognostic models.

Emerging evidence suggests that gliomas in populations underrepresented in research may follow distinct biological and clinical trajectories. Recently, several studies have reported that Black and Hispanic patients (AMR) with GBM have equal or better survival than non-Hispanic White patients, even after adjustment for age, treatment, and comorbidities [9–14]. However, it remains unclear whether the germline risk alleles and somatic mutation patterns are equally relevant, or occur at similar frequencies, across these ancestral backgrounds.

To address these gaps, we assembled a multicenter research team and developed a multi-institutional biospecimen and data resource that integrates clinically annotated glioma tissues from patients with enriched representation of African (AFR), Admixed American (AMR), and Asian (AS) ancestries. This paper describes the design and development of this resource, summarizes demographic and molecular features across ancestral subgroups, and illustrates how this cohort can be used for multi-ancestry analyses in neuro-oncology. Specifically, we describe whole exome sequencing (WES) data and associated clinical information across participating institutions. By establishing this foundation, we aim to support more inclusive biomarker discovery, improve risk prediction, and advance precision-medicine approaches that better reflect the full genetic and clinical diversity of glioma (Figure 1).

**Figure 1.**
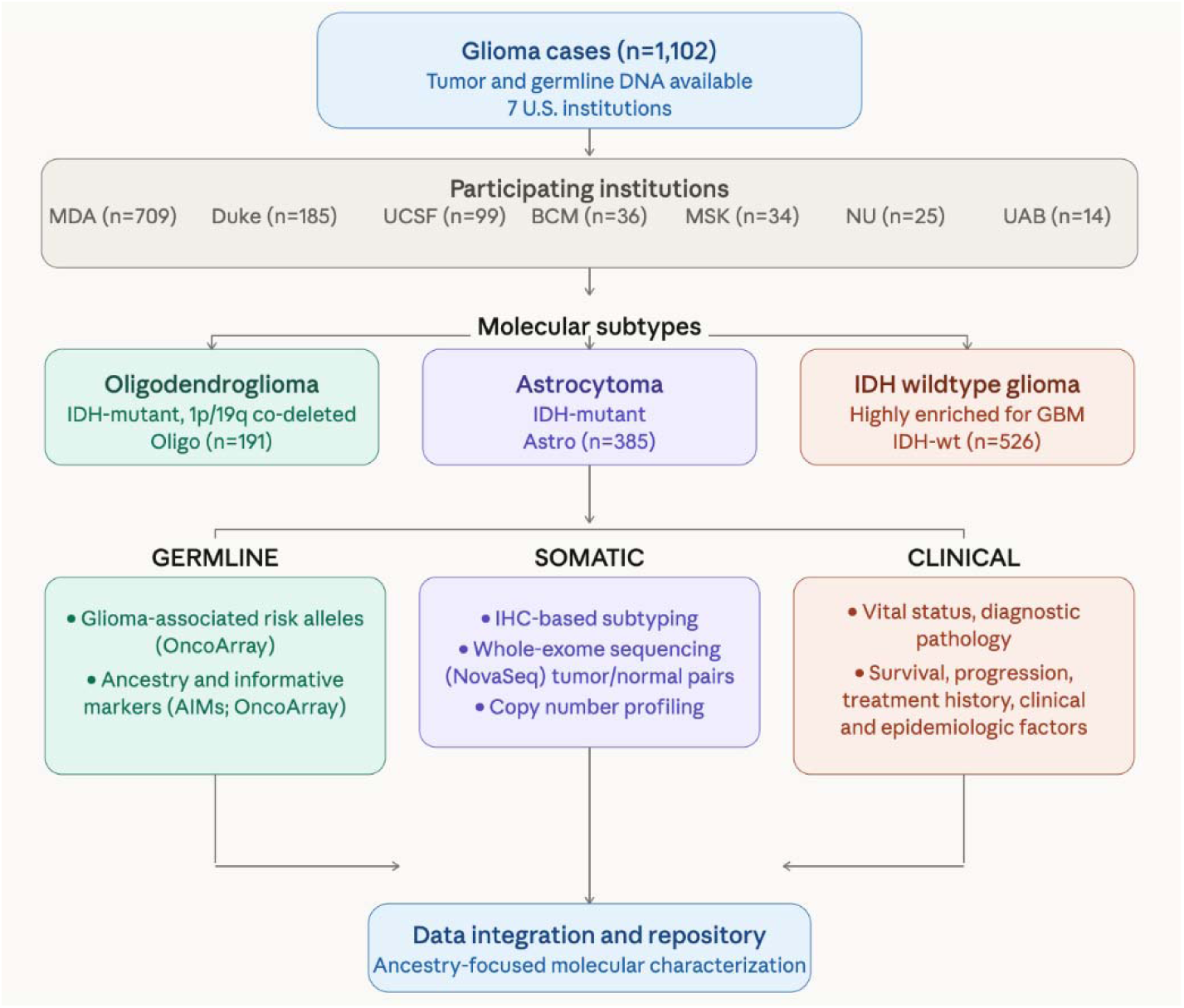
Overview of the study design and analytical framework. Schematic representation of the multi-institutional glioma study workflow (7 U.S institutions), illustrating cohort assembly (n=1,102), data integration, and downstream analyses. Clinical and demographic information, together with harmonized somatic and germline genomic data, are integrated across participating sites to enable ancestry-focused characterization of molecular alterations and mutational processes across glioma tumor types.

## METHODS

### Study Cohort and Sample Collection

Tumor and blood specimens were obtained from patients who had participated in the Glioma International Case-Control (GICC) study [19]. In addition, we invited investigators at other centers including those supported by the NCI-funded Brain Tumor SPORE program, to collect samples and increase the diversity of the cohort, since some of those centers service primarily underrepresented populations. Primary collections with patient data and tissue samples originated from MD Anderson Cancer Center (1985-2023), Baylor College of Medicine (2004-2021), Duke University (1983-2023), and Memorial Sloan Kettering Cancer Center (MSK) (2010-2013). All contributing sites obtained approval from their respective Institutional Review Boards (IRBs), and written informed consent was obtained from all participants at the time of sample collection.

### Clinical and Demographic Data Abstraction

Demographic and clinical data, including age at diagnosis, sex, self-reported race/ethnicity, clinical presentation, treatment, and vital status, were abstracted from medical records at each institution using standardized data forms and then harmonized across sites.

### Genomic DNA Extraction and Quality Control

All tumor tissue specimens were processed centrally for consistency. Tumor DNA was extracted from representative formalin-fixed paraffin-embedded (FFPE) tissue blocks in Dr. Huse’s laboratory at MD Anderson Cancer Center using a Purigen device and reagents, according to manufacturer’s instructions (Purigen Biosystems). Tumor and germline DNA was quantified via spectrophotometry and fluorometry and assessed for integrity by agarose gel electrophoresis. Aliquots of each sample were stored at -80°C until shipment to the Human Genome Sequencing Center at Baylor College of Medicine.

### Sequencing and Genotyping

WES was performed on all samples meeting DNA quality criteria using UV/Vis spectrometry and “yield gels” to estimate DNA sample integrity. WES libraries were prepared using KAPA Hyper reagent and 500 ng of gDNA. A set of ∼96 unique dual indexed barcoded adaptors from Illumina (cat # 20022370) were used to multiplex samples [20]. For WES enrichment, 10 pre-capture libraries were pooled and enriched by hybridizing to either the HGSC-VCRome 2.1 with spike in probes (Roche NimbleGen custom design) or HGSC-ClinExD probes (Twist custom design) [21]. High-throughput sequencing was performed using the Illumina NovaSeq 6000 as a 70-plex per S4 flow cell lane to achieve ∼100x average coverage. For cases not previously genotyped within the parent GICC study [20], genome-wide genotyping was performed using the Illumina Infinium Global Diversity Array (GDA) according to manufacturer’s instructions. SNPs were analyzed using GenomeStudio Software (Version 2011.1, Illumina Inc.)

### Bioinformatic Processing and Variant Calling

Raw sequencing reads underwent stringent quality control using FastQC and MultiQC, and samples or libraries failing minimum thresholds for base quality, coverage, or duplication rates were excluded from further analysis. Reads were aligned to the hg38 human reference genome using BWA-MEM v0.7.17, followed by duplicate removal with Sambamba v0.8.1 [22] and base quality score recalibration with GATK v4.3.0.0.[23] Somatic variants were called using MuSE v2 [24] and annotated with Ensembl Variant Effect Predictor (VEP). Additional quality control included cross-checks for sample mix-ups, review of batch effects, and confirmation of ancestry assignments using principal component analysis against reference populations. Germline WES data were used for ancestry assignment via KING [25]. Copy Number Aberrations (CNA) were computed using Sequenza v3.0.0 [26]. Genes were classified as amplified if total copy number exceeded ploidy + 5, or deleted if copy number was at or below ploidy - 2.

### Ancestry and Sex Inference

Genetic ancestry inference was performed using KING, based on germline variants (processed from WES data) obtained from GATK HaplotypeCaller (v4.3.0). Variants were first called for each normal (germline) sample on the GRCh38 reference genome and then lifted over to hg19 coordinates using the LiftOver function in R to ensure compatibility with population reference panels. The lifted variant files were converted to PLINK binary format (BED/BIM/FAM) using plink2. rsID annotations were added to BIM files in R to facilitate SNP matching with the 1000 Genomes reference dataset, The resulting SNP panels were used as input to KING to estimate genetic ancestry by comparing each sample to reference populations from the 1000 Genomes Project. Ancestry was assigned as per Anc_1^st^ on KING’s output, which is based on the maximum probability, hence no thresholds were required.

Sex was inferred from germline WES BAM files for cases with missing gender data using chromosome-specific alignment statistics. For each sample, alignment counts were extracted with samtools idxstats. For chromosomes X and Y, we calculated the ratio of mapped reads on chromosome X (Y:X read ratio). A threshold of 0.025 for the Y:X ratio was used to infer chromosomal sex, with values below this threshold classified as female and values at or above this threshold classified as male. Reported and inferred sex were compared for all samples; one discordant sample was excluded from further analysis.

### Mutational Signature Fitting

Mutation count matrices (96 trinucleotide context) were generated from somatic VCF files using the hg38 reference genome. The R package MutationalPatterns [27] was used to fit COSMIC mutational signatures to each samples profile. Mutational signature analysis used COSMIC single base substitution (SBS) signatures version 3.2 as the reference set.Mutational signatures were fitted using the strict refitting approach with iterative backwards selection (max_delta = 0.004) to remove signatures that did not contribute significantly to the observed mutation profiles. This approach reduces overfitting by retaining only signatures that explain a substantial proportion of the sample-specific mutation spectrum.

### Survival Analysis

Overall survival was analyzed using the Kaplan–Meier methods, stratified by institution/site. IBM SPSS Statistics (version 29.0.2.0; RRID: SCR_002865) was used for these analyses, and log-rank p-values < 0.05 were considered statistically significant. IBM SPSS Statistics software was also used to perform univariate and multivariate survival analysis with Cox proportional hazards model and a 2-sided *p*-value of less than 0.05 was considered statistically significant. OS defined from date of diagnosis to death or last follow-up.

### Statistical Analysis

To summarize demographic and clinical characteristics, Chi-square and Fisher’s exact tests were applied to compare categorical variables, and t-tests or ANOVA were used for continuous variables, as appropriate. Associations between mutational signature contributions and tumor type were assessed using the Kruskal-Wallis test, a non-parametric test for comparing distributions across multiple groups. P-values were adjusted for multiple comparisons using the Benjamini-Hochberg false discovery rate method (FDR < 0.05). All statistical analyses were conducted in R (version 4.2).

## RESULTS

### Patient Demographics and Molecular Subtypes

The cohort included 1,102 glioma patients from seven institutions (Duke n=185, MDA n=709, MSK n=34, BCM n=36, UCSF n=99, UAB n=14, and NU n=25) classified into three molecular subtypes: Oligodendroglioma, IDH-mutant and 1p/19q co-deleted (Oligo; n=191), astrocytoma, IDH-mutant (Astro, n=385), and IDH wildtype gliomas, highly enriched for GBM (IDH-wild type; n=526) (Table 1). European (EUR) ancestry was most prevalent (66.9%), followed by Admixed American (AMR) at 21.1%, African (AFR) at 10.3%, and Asian (AS) at 1.6% (Table 1). Principal Components Analysis (PCA) of germline ancestry data showed distinct clusters as expected (Supplementary Figure 1) Due to the lower number of AS cases in the cohort, they have been included in analysis only where statistically possible. The overall inferred sex distribution showed a modest male predominance (57.5%), with males comprising 51.8% of Oligo, 58.2% of Astro, and 59.1% of IDH-wt cases. Patients with IDH-mutant tumors were diagnosed at significantly younger median ages than those with IDH-wild type tumors (p<0.001), with mean ages of 41.1 years for Oligo and 35.3 years for Astro, compared with 56.1 years for IDH-wt gliomas (Table 1).

**Table 1.**
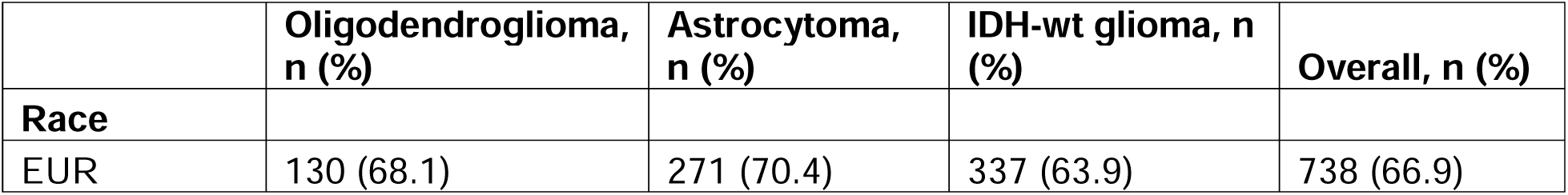
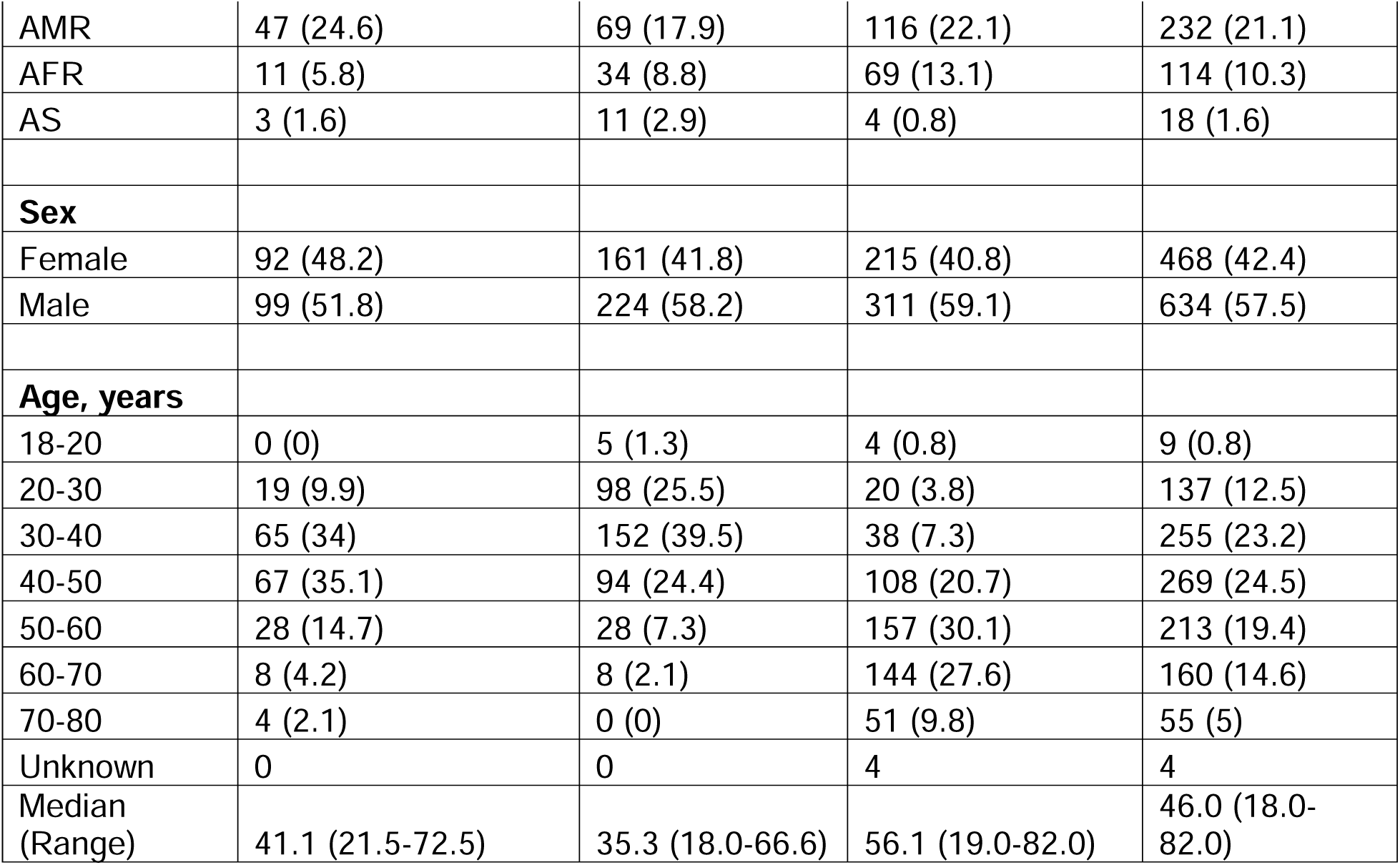
Characteristics and molecular subtype-specific of the multi-institutional cohort.

|  | Oligodendroglioma, n (%) | Astrocytoma, n (%) | IDH-wt glioma, n (%) | Overall, n (%) |
| --- | --- | --- | --- | --- |
| **Race** |  |  |  |  |
| EUR | 130 (68.1) | 271 (70.4) | 337 (63.9) | 738 (66.9) |
| AMR | 47 (24.6) | 69 (17.9) | 116 (22.1) | 232 (21.1) |
| AFR | 11 (5.8) | 34 (8.8) | 69 (13.1) | 114 (10.3) |
| AS | 3 (1.6) | 11 (2.9) | 4 (0.8) | 18 (1.6) |
| **Sex** |  |  |  |  |
| Female | 92 (48.2) | 161 (41.8) | 215 (40.8) | 468 (42.4) |
| Male | 99 (51.8) | 224 (58.2) | 311 (59.1) | 634 (57.5) |
| **Age, years** |  |  |  |  |
| 18-20 | 0 (0) | 5 (1.3) | 4 (0.8) | 9 (0.8) |
| 20-30 | 19 (9.9) | 98 (25.5) | 20 (3.8) | 137 (12.5) |
| 30-40 | 65 (34) | 152 (39.5) | 38 (7.3) | 255 (23.2) |
| 40-50 | 67 (35.1) | 94 (24.4) | 108 (20.7) | 269 (24.5) |
| 50-60 | 28 (14.7) | 28 (7.3) | 157 (30.1) | 213 (19.4) |
| 60-70 | 8 (4.2) | 8 (2.1) | 144 (27.6) | 160 (14.6) |
| 70-80 | 4 (2.1) | 0 (0) | 51 (9.8) | 55 (5) |
| Unknown | 0 | 0 | 4 | 4 |
| Median<br>(Range) | 41.1 (21.5-72.5) | 35.3 (18.0-66.6) | 56.1 (19.0-82.0) | 46.0 (18.0-82.0) |

### Survival Outcomes

Median overall survival (OS) differed markedly by molecular subtype, consistent with multiple prior reports [3, 16–18]. Estimate median OS was 15.7 years (95% CI, 14.1–19.2) for Oligo, 10.6 years (95% CI, 9.2–13.4) for Astro, and 1.9 years (95% CI, 1.8–2.1) for IDH-wt glioma. Survival analyses were conducted as described in Methods, with OS defined from date of diagnosis to death or last follow-up.

We compared the OS within each subtype (Fig.2A) and ancestry (Fig.2B-D). Consistent with prior literature [3, 16–18], significant survival differences were observed for IDH-mut compared with IDH-wt tumors (p<0.001). IDH-wt glioma showed significantly worse survival than Oligo (HR [95% CI]: 8.1 [6.3–10.1], p<0.001) and Astro (HR [95% CI]: 5.1 [4.3–6.1], p<0.001). When comparing survival difference within IDH-mut gliomas, Oligo expectedly exhibited significantly improved survival vs Astro (HR [95% CI]: 0.63 [0.48–0.83], p<0.001; Fig.2A). The difference in overall survival across subtypes remained significant after adjusting for age, gender, and institution in multivariate analysis (p<0.001).

**Figure 2.**
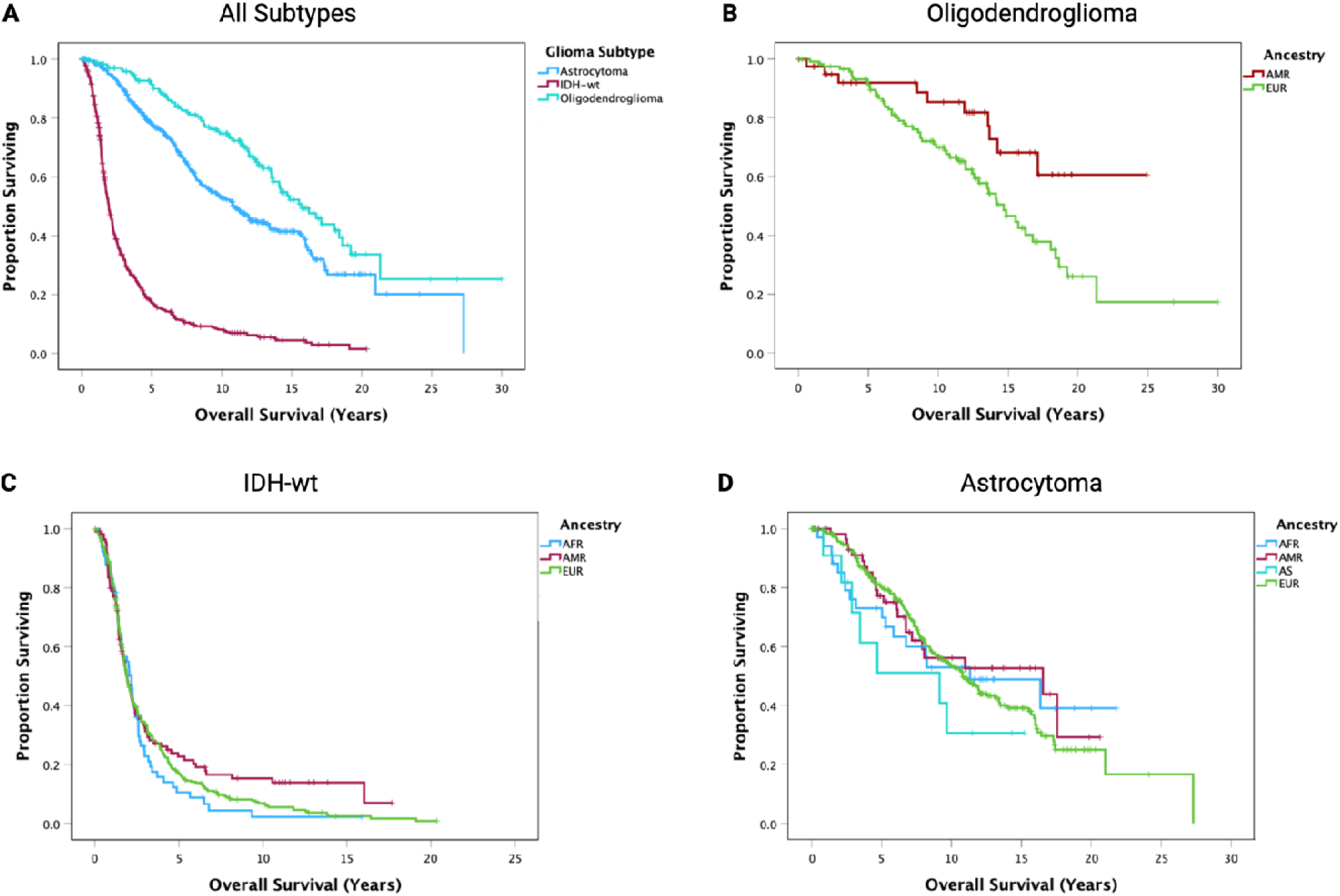
Overall survival across glioma subtypes and ancestry groups. Kaplan–Meier curves comparing overall survival (OS) by glioma subtype and genetic ancestry. (A) OS across all glioma subtypes demonstrates significant survival differences between IDH-mut and IDH-wt tumors (p<0.001). Oligo vs Astro: HR 0.63, 95% CI 0.48–0.83; p<0.001. IDH-wt vs Oligo: HR 8.1, 95% CI 6.3–10.1; p<0.001. IDH-wt vs Astro: HR 5.1, 95% CI 4.3–6.1; p<0.001. (B) Within oligo, the AMR ancestry subgroup demonstrated significantly improved survival compared with EUR (HR 0.67, 95% CI 0.48–0.94; p=0.011). (C) In IDH-wt tumors (D) In Astro. Ancestry subgroups with fewer than five events were excluded from subgroup analyses. Multivariate analysis was performed to adjust for age, sex and site. Tick marks indicate censored observations. Log-rank p-value<0.01 was considered statistically significant. AMR: Admixed American; AFR: African; AS: Asian; EUR: European.

Comparing different ancestry groups within Oligo, significantly improved survival was observed for AMR subgroup compared with EUR (Fig.2B; HR [95% CI]: 0.67 [0.48–0.94], p=0.011). The survival difference between AMR and EUR subgroups remained statistically significant after adjustment for clinical covariates in multivariable analysis (adjusted for age, sex and site). In IDH-wt tumors (Fig.2C), no significant survival difference was observed between the ancestry groups. However, a trend for improved survival in the AMR group when compared with the AFR group was noted (p=0.16). Similarly, in Astro (Fig.2D), no significant survival difference was observed between the ancestry group, but an improved survival trend in the AMR group relative to the AS group was noted (p=0.11). Specific ancestry groups were omitted from analysis where the number of events in each subgroup was less than 5.

### Somatic alterations across the sample cohort

WES data were used to identify mutations and copy-number alterations (FIG. 3A-3B). Overall, the observed patterns were consistent with established genomic profiles of Astro, Oligo, and IDH-wt glioma. Astros showed high frequencies of mutations in *TP53*, and ATRX, Oligos were characterized by TERT promoter mutations, and showed enrichment for mutations in CIC, FUBP1, and NOTCH1. IDH-wt gliomas demonstrated frequent mutations and/or copy number alterations involving EGFR, PTEN, TP53, NF1, CDK4, MDM2, and CDKN2A/B, along with TERT promoter mutations.

**Figure 3.**
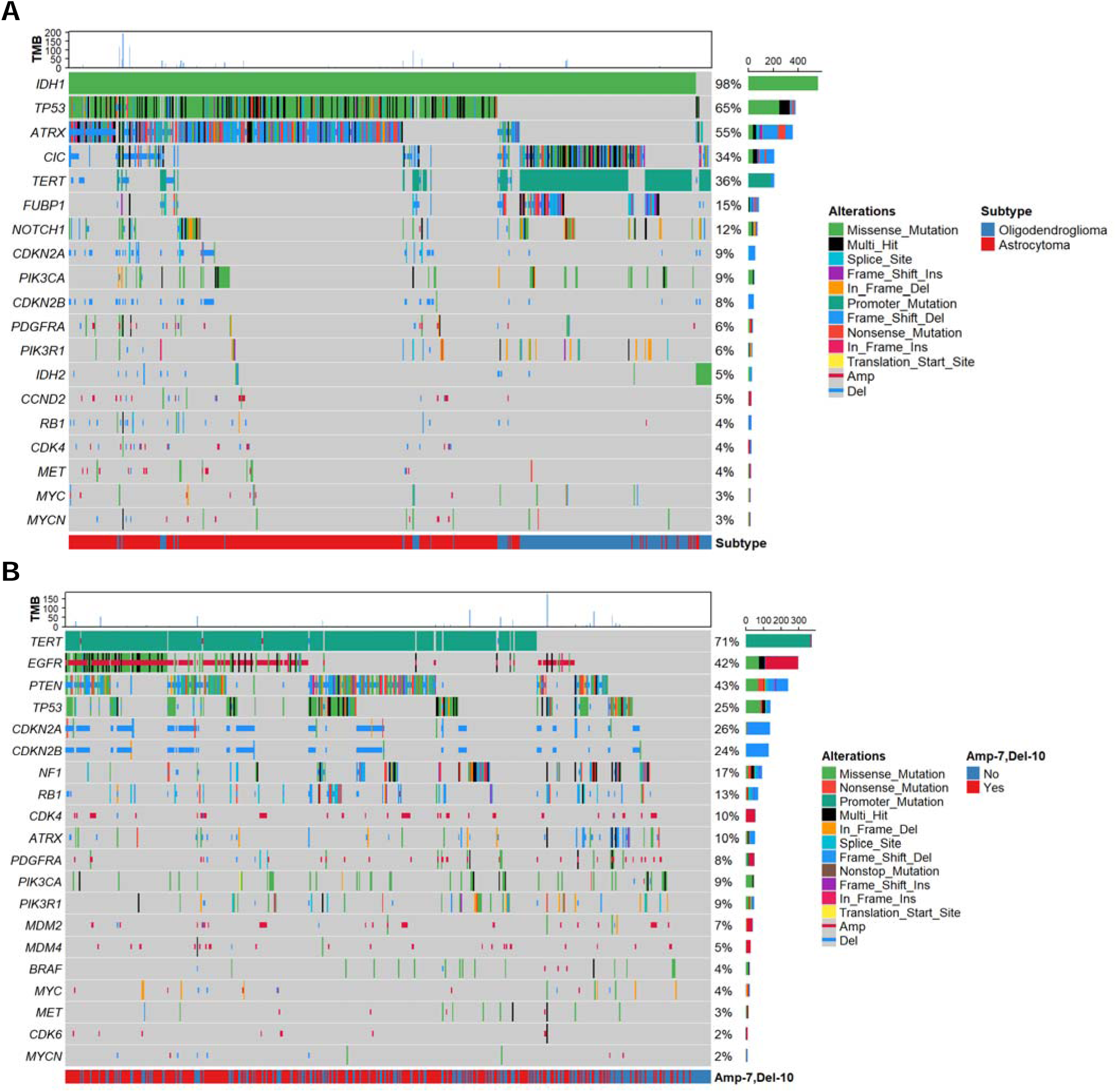
Oncoplots of recurrent somatic alterations across the glioma cohort. Oncoplots summarize somatic mutations and copy-number alterations across (A) 576 IDH-mutant and (B) 526 IDH-wt glioma samples. Rows represent genes recurrently altered in the cohort, and columns represent individual tumors. Alteration types are color-coded and include missense, nonsense, frameshift, splice-site, in-frame insertions/deletions, promoter mutations, and copy number gains or losses, as indicated in the legend. The bar plot on the right shows the percentage of samples altered for each gene, while the bar plot at the top indicates the total number of alterations per sample. Tumor subtype annotations at the bottom distinguish Astro from Oligo, and IDH-wt glioma with and without combined gain of chromosome 7 and loss of chromosome 10 (+7/-10).

Most genes exhibited similar alteration frequencies following stratification by ancestral group (FIG. 4A-B). Notably, the tumor suppressor ATRX, which is commonly inactivated in Astros, showed significantly higher rates of alteration in AMR and AFR patients with IDH-wt glioma (20% and 16%, respectively), compared with EUR patients (6%) (Fig.4A; p < 0.001 and p = 0.004; two sample proportion tests for AMR vs EUR and AFR vs EUR, respectively). Moreover, in IDH-wt glioma, AMR patients had significantly lower frequencies of EGFR alterations (32%) compared with AFR and EUR patients (45–51%; p<0.05; Fig. 4B). These findings suggest that, while overall genomic patterns are broadly shared, specific alterations may differ by ancestry within subtypes.

**Figure 4.**
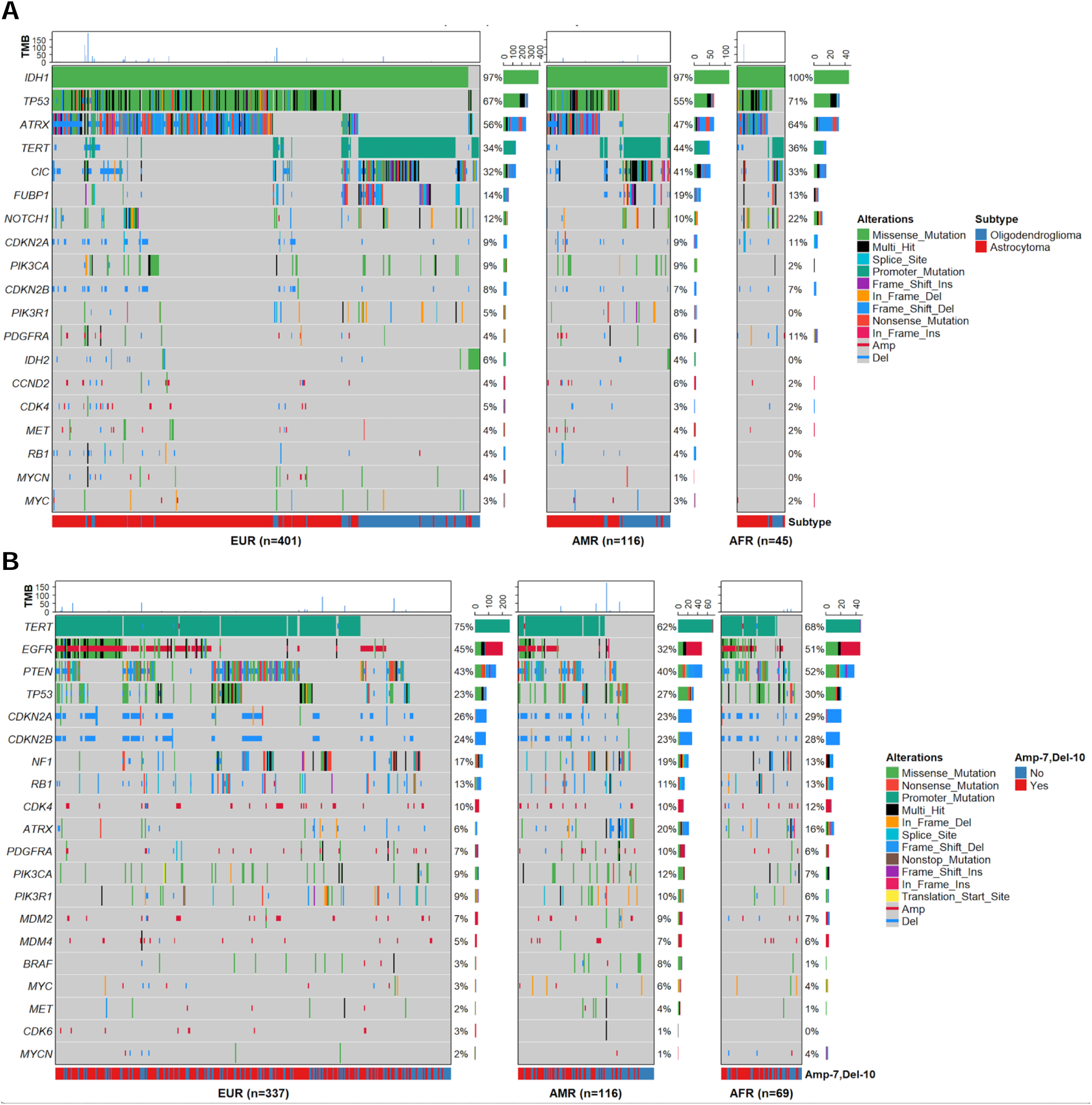
Ancestry-stratified oncoplots of recurrent somatic alterations in diffuse glioma. Oncoplots summarize somatic mutations and copy-number alterations across glioma tumors stratified by genetic ancestry and molecular subtype. Panels show (A) IDH-mutant and (B) IDH-wt gliomas for individuals of AFR, AMR, and EUR ancestry. Alteration types are color-coded and include missense, nonsense, frameshift, splice-site, in-frame insertions/deletions, promoter mutations, and copy number gains or losses, as indicated in the legend. The percentages of tumors with alterations in each gene within the corresponding ancestry group are indicated. Tumor subtype annotations at the bottom of IDH-mut group distinguish Astro from Oligo, and IDH-wt gliomas with and without combined gain of chromosome 7 and loss of chromosome 10 (+7/−10).

Further subgroup-specific survival analyses revealed that ATRX alterations were associated with significantly improved overall survival in the AFR subgroup (p = 0.003), while no significant survival association was observed in AMR or EUR patients (Supplementary Figure. 2). In contrast, analysis of EGFR alterations showed no significant associations with survival across any ancestry group (data not shown).

### Glioma subtypes exhibit distinct and heterogeneous mutational signature profiles

Single-nucleotide variant (SNV) data from 1,102 glioma patients were analyzed to characterize underlying mutational processes across tumor types. To obtain robust and biologically meaningful assignments, we applied a strict signature-fitting procedure that iteratively removed signatures not improving reconstruction of the observed mutational profiles, thereby reducing overfitting and minimizing spurious contributions. This approach identified a subset of SBS signatures consistently reflected in the mutational spectra in this cohort.

We tested whether these specific mutational signatures were associated with glioma tumor type. Eighteen SBS signatures showed statistically significant differences in contribution across glioma subtypes (FDR < 0.05), indicating tumor-type specific mutational processes (Supplementary Table 1). These signatures span several etiologic categories, including DNA repair deficiency–associated processes, therapy-associated mutational processes, environmental or exogenous exposures, and age-related processes, while some have unknown or uncertain etiologies (for example SBS23, SBS37, SBS19, and SBS94) (Supplementary Table 1).

To explore broader patterns of mutational processes across glioma subtypes, we calculated the mean contribution for each signature within each tumor type and visualized these patterns using hierarchical clustering (Supplementary Figure 3). This process revealed signatures with shared enrichment across subtypes (for example SBS31, SBS37, SBS94) as well as signatures more specifically enriched in particular groups, including SBS11, SBS37, SBS7b, and SBS1 in Oligo; SBS87, SBS94, SBS23, SBS19, SBS86, SBS32, and SBS6 in Astro; and SBS1, SBS40, SBS38, SBS4, SBS44, SBS3, and SBS8 in IDH-wt glioma (Supplementary Table 1).

We next focused on signatures identified as statistically significant in tumor-type association analysis, assessing their mean contributions by heat map (Figure 5A-B). This analysis identified mutational processes that distinguish glioma subtypes (SBS37, SBS94, SBS1, SBS87, SBS23, SBS19, SBS6, and SBS11). Three out of eight signatures are associated with therapy response (SBS11, SBS87 and SBS94 while others are aging-related (SBS1), DNA repair deficiency-related (SBS6), environmental (SBS37), or of unknown etiology (SBS19 and SBS23). Astro was significantly associated with SBS94, SBS87, SBS23, SBS19 and SBS6, Oligo was significantly associated with SBS37 and SBS11, and IDH-wt glioma was significantly associated with SBS1.

**Figure 5.**
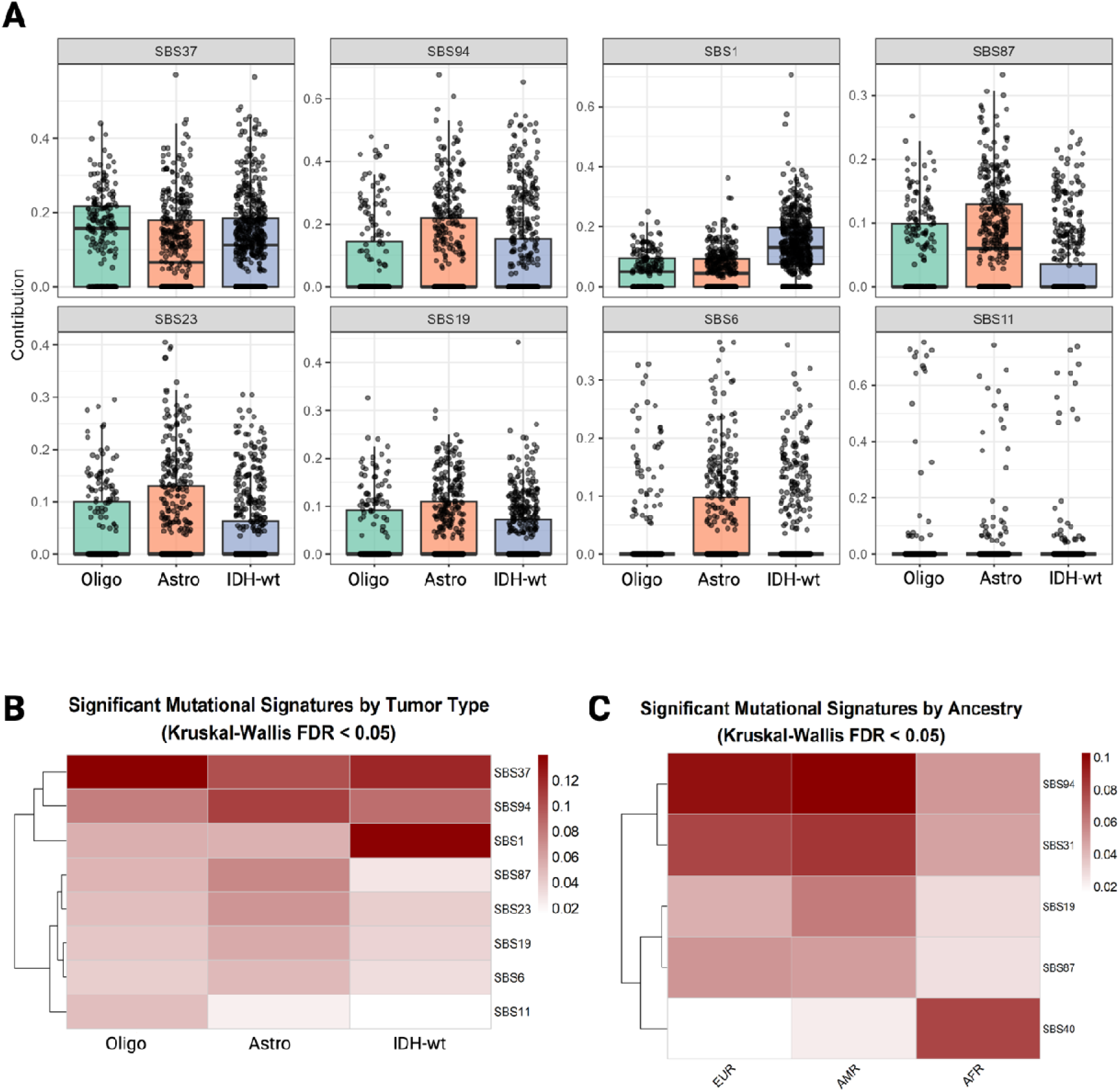
Tumor type and ancestry–associated mutational signature contributions in glioma. (A) Boxplots showing COSMIC single base substitution (SBS) mutational signature contributions stratified by glioma subtype (Oligo, Astro, and IDH-wt glioma). Only signatures demonstrating statistically significant differences across tumor types by Kruskal–Wallis testing (FDR < 0.05) are displayed. Points represent individual tumors, boxes indicate interquartile ranges with medians, and adjusted p-values are shown for each signature. Proposed biological etiologies (for example, temozolomide exposure, defective DNA mismatch repair, ultrav olet light–associated processes) are annotated based on COSMIC definitions. Together, these boxplots illustrate that specific mutational processes are differentially active across glioma subtypes. (B) Heatmap summarizing tumor type and (C) ancestry–associated SBS signatures identified by Kruskal–Wallis testing (FDR < 0.05). Rows represent significant mutational signatures and columns represent tumor types/ancestry groups. Color intensity reflects the mean signature contribution within each subtype. Hierarchical clustering highlights differences in relative signature activity and emphasizes mutational processes that distinguish glioma molecular subgroups.

Using the same approach, we identified ancestry-associated mutational signatures (Fig. 5C). SBS94, SBS31, SBS19, SBS87 and SBS40 signatures showed statistically significant association ancestry groups. While etiologies for SBS94, SBS19, and SBS40 are “unknown”, SBS31 is a platinum chemotherapy associated signature, and SBS87 is a Thiopurine chemotherapy associated signature. These two chemotherapy-related signatures were significantly enriched in EUR and AMR ancestry groups compared with the AFR group (Fig.5C).

Finally, we extended this analysis to the individual sample level, demonstrating statistically significant signature contributions across all 1,102 tumors, thereby capturing intra-tumor type heterogeneity and highlighting variability in mutational processes among patients (Supplementary Figure. 4). Collectively, these analyses show that glioma subtypes are characterized by distinct, biologically interpretable mutational signatures, while also revealing substantial heterogeneity within tumor classes.

## DISCUSSION

This large, multi-institutional cohort integrates harmonized clinical, genomic, and ancestry data from 1,102 adults with diffuse glioma to define how germline ancestry, molecular subtype, and mutational processes jointly shape tumor biology and outcomes. To our knowledge, this represents the largest ancestrally diverse cohort of glioma patients with detailed clinical data, integrated molecular and mutational signature profiling reported to date. This cohort enables detailed characterization of ancestry-associated variation in glioma, supporting integrated analyses of survival, somatic alterations, and mutational signatures across IDH-mutant and IDH-wildtype subtypes.

### Demographic Distribution

Our findings regarding ancestry distribution, demographic characteristics, and survival outcomes in this cohort are broadly consistent with prior epidemiologic and clinical studies [12, 28, 29]. For instance, we observed an overall male predominance in our cohort across ancestries (male-to-female ratios: whole cohort ∼1.3:1; EUR 1.4:1; AMR 1.2:1; AFR 1.3:1; AS 1.3:1), with no significant difference between groups (Fisher’s exact test p=0.59). Prior population-based studies and clinical series have demonstrated that males are more susceptible to gliomas across molecular subtypes and that IDH-mutant gliomas tend to present in younger adults, whereas IDH-wt tumor, are more common in older individuals [9, 28]. Although the magnitude of male preponderance that we observed is somewhat lower than that reported in some largely European-ancestry cohorts possibly due to the large proportion of lower grade gliomas in the cohort, it remains directionally concordant with the established sex ratio in glioma [29]. Moreover, IDH-mutant subtypes had younger age at onset, consistent with previously reported epidemiological patterns [3, 16–18, 30]. Which is consistent with prior epidemiologic evidence [30]. Taken together, these characteristic patterns confirm that this cohort is fundamentally similar to nationwide glioma characteristics across molecular tumor subtype, age and gender [17, 18, 31].

Survival outcomes stratified by molecular subtype we observed were also similar to contemporary reports from TCGA and other datasets [17, 18, 31]. We found markedly longer median survival for Oligos and Astros (medians of 15.7 and 10.6 years, respectively, compared with 1.9 years for IDH-wt gliomas). These survival differences align with the prognostic hierarchy established in TCGA analyses and other seminal studies demonstrating the substantial outcome advantage conferred by IDH mutation and 1p/19q codeletion [3, 16–18]. Our cohort further confirms the aggressive nature and correspondingly poor prognosis of adult IDH-wt glioma across institutions.

Within subtype-stratified analyses, we observed ancestry-associated survival differences that were most pronounced in Oligo. Specifically, patients of AMR ancestry demonstrated significantly improved survival compared with EUR patients, and this association remained robust after adjustment for age, sex, and institution. In contrast, no statistically significant ancestry-associated survival differences were observed in IDH-wt or Astro, although trends toward improved survival in the AMR subgroup were noted in both settings. Our findings are consistent with prior population-based studies reporting improved survival among AMR populations in glioma, including Oligo, supporting the observation that AMR ancestry confers a survival advantage for Oligo patients in our cohort [9, 32–34]. These findings suggest that ancestry-related survival differences may be subtype-specific and potentially more detectable in biologically favorable tumor contexts such as Oligo. The observed patterns raise the possibility that genetic background, tumor biology, or differential treatment response may contribute to survival heterogeneity within molecularly defined glioma subtypes.

### Ancestry, incidence, and outcomes

Differences in glioma incidence and outcomes across racial and ethnic groups likely arise from a complex interplay of inherited genetic susceptibility, tumor intrinsic biology, and non-biologic factors. Germline studies have shown that individuals of European descent are enriched for glioma risk alleles, contributing to higher incidence in this group relative to non-European ancestry populations [35]. Moreover, increasing European genetic ancestry in admixed groups Hispanic/Latino populations has been associated with elevated glioma risk, supporting a dose–response relationship between ancestry proportion and susceptibility [9]. Somatic alterations also appear to vary by ancestry, with prior work suggesting differences in the frequency spectrum of mutations across key glioma pathways, including TP53, ATRX, and receptor tyrosine kinase networks. Our cohort, which included both AFR and AMR ancestries, is consistent with these findings and provides a platform for future analyses that jointly model genetic ancestry, germline risk alleles, and outcomes [36]. For example, we observed increased rates of ATRX mutations in AFR and AMR ancestry groups relative to EUR counterparts in IDH-wt gliomas, further supporting ancestry-associated differences in tumor biology [35]. Subgroup survival analysis revealed an improved survival in AFR patients with ATRX alterations, suggesting that the prognostic impact of genomic alterations may be context-dependent and vary across ancestral backgrounds, underscoring the importance of ancestry-aware biomarker evaluation.

Our results confirm prior findings that differences in survival between racial and ethnic groups are multifactorial, involving complex genetic, molecular, and social determinants. Our cohort’s pattern of survival differences between ancestry groupings concurs with published data and further establishes the need to broaden representation across different populations in glioma genomic and clinical research.

### Mutational Signatures

Mutational signature analysis provided additional insight into the biological, therapeutic processes shaping glioma genomes in this cohort [37]. By focusing on signatures significantly associated with tumor type and ancestry group, we identified distinct mutational profiles across Oligo, Astro, and IDH-wt gliomas, and across EUR, AMR and AFR ancestry groups, reinforcing and extending prior work that has largely examined smaller or less ancestrally inclusive series. Our results demonstrate the importance of considering the joint roles of established factors such as of age- and therapy-associated processes in adult glioma along with the less well-characterized signatures [38].

A key finding is the prominence of therapy-related signatures, particularly SBS11 and SBS87. SBS11, the canonical temozolomide signature, was enriched in treated IDH-mutant tumors, in line with previous longitudinal and recurrence-focused studies showing temozolomide-driven hypermutation and signature emergence [37, 39]. The additional enrichment of SBS87 in specific IDH-mutant subgroups suggests that alkylating chemotherapy and related agents may leave a more complex mutational footprint in glioma than SBS11 alone, an observation that has been less consistently emphasized in prior glioma-specific reports [38, 39]. Chemotherapy-related signatures SBS87 and SBS31 were also significantly different among ancestry groups, showing particular enrichment in EUR and AMR groups compared with AFR. These results support the hypothesis that therapy-related mutational processes contribute substantially to the evolutionary landscape of glioma and likely vary across both molecular subtypes and patient ancestry groups. These findings demonstrate again the importance of considering treatment exposure and population diversity when interpreting mutational signatures and their potential clinical implications.

Age-related and DNA repair–related signatures showed patterns broadly concordant with the existing literature but with added resolution by molecular subtype [40]. SBS1, reflecting clock-like, age-associated mutagenesis, was detectable across all groups but was most prominent in IDH-wt glioma patients, paralleling their older age at diagnosis and aligning with prior pan-cancer and CNS-focused signature analyses [37]. Enrichment of SBS6 and related mismatch repair–associated signatures in a subset of Astros is consistent with previous descriptions of MMR-deficient or hypermutated gliomas, often in the setting of prior alkylator therapy or germline/somatic MMR pathway alterations [41]. Together, these findings reinforce that age and DNA repair capacity remain major determinants of the mutational milieu in adult diffuse glioma.

Importantly, we also observed robust contributions from signatures with uncertain or incompletely defined etiology, including SBS19, SBS23, and SBS37, particularly within Astros. Prior studies have variably detected these signatures in glioma or pan-cancer datasets but have not consistently linked them to specific molecular subtypes or ancestrally diverse populations [38]. The enrichment of these signatures in our cohort suggests that additional, as yet uncharacterized mutational processes potentially related to endogenous metabolism, environmental exposures, or context-specific therapeutic effects may operate in distinct glioma subsets. The association of SBS23 with more aggressive behavior in other series, together with its prominence in our Astro group, raises the possibility that certain “unknown” signatures could serve as markers of biological aggressiveness or treatment history once better understood.

### Implications and Future Directions

Taken together, our mutational signature findings show strong concordance with prior work regarding the central roles of temozolomide exposure, age, and mismatch repair status, while adding new detail about the distribution of therapy-related and unknown signatures across modern WHO-defined subtypes in an ancestrally diverse cohort. Future analyses within this resource could integrate signatures with treatment data, and longitudinal outcomes to test whether specific signatures predict response, recurrence patterns, or survival within and across ancestry groups. Such work could clarify the biological basis of currently unassigned signatures, refine risk stratification, and ultimately inform ancestry-aware, exposure-aware precision strategies in glioma.

## Ethics Statement

This study was conducted in accordance with the ethical principles outlined in the Declaration of Helsinki and all applicable federal and institutional regulations. Collection and analysis of clinical, genomic, and demographic data were approved by the Institutional Review Boards (IRBs) of all participating institutions. Written informed consent was obtained from all participants or their legally authorized representatives, with consent permitting genomic sequencing, data harmonization, and secondary research use.

## Authorship

**Conceptualization**: JTH, JB

**Study design**: MB, JTH, CIA, JB, QTO

**Data acquisition and interpretation**: MB, JTH, CIA, JB, BP, QTO, KMW, VK, AD, MWr, MWo, GA, BN, LN, JR, AP, VT, JM, JH, HD, XL, ZM, PM

**Sample processing**: IO, TB

**Formal analysis**: ST, HN, JTH, KF

**Writing-original draft**: MB, HN, ST, JTH, CIA, JB

**Writing-review and editing**: CIA, JTH, JB, CH, DMM, HN

**Supervision**: MB, CIA, JTH, JB

## Data Availability

The data will be made available upon request to the corresponding author and will be subjected to approval by the relevant institutional review boards, data-use agreements, and applicable regulations.

## References

1. Louis, D.N., A. Perry, P. Wesseling, D.J. Brat, I.A. Cree, D. Figarella-Branger, et al., The 2021 WHO Classification of Tumors of the Central Nervous System: a summary. Neuro Oncol, 2021. 23(8): p. 1231–1251.

2. Price, M., C.A.P. Ballard, J.R. Benedetti, C. Kruchko, J.S. Barnholtz-Sloan, and Q.T. Ostrom, CBTRUS Statistical Report: Primary Brain and Other Central Nervous System Tumors Diagnosed in the United States in 2018-2022. Neuro Oncol, 2025. 27(Supplement_4): p. iv1–iv66.

3. Brennan, C.W., R.G. Verhaak, A. McKenna, B. Campos, H. Noushmehr, S.R. Salama, et al., The somatic genomic landscape of glioblastoma. Cell, 2013. 155(2): p. 462–77.

4. Tamimi, A.F. and M. Juweid, *Epidemiology and Outcome of Glioblastoma*, in Glioblastoma, S. De Vleeschouwer, Editor. 2017: Brisbane (AU).

5. Kinnersley, B., J.S. Mitchell, K. Gousias, J. Schramm, A. Idbaih, M. Labussiere, et al., Quantifying the heritability of glioma using genome-wide complex trait analysis. Sci Rep, 2015. 5: p. 17267.

6. Stupp, R., W.P. Mason, M.J. van den Bent, M. Weller, B. Fisher, M.J. Taphoorn, et al., Radiotherapy plus concomitant and adjuvant temozolomide for glioblastoma. N Engl J Med, 2005. 352(10): p. 987–96.

7. Buckner, J.C., E.G. Shaw, S.L. Pugh, A. Chakravarti, M.R. Gilbert, G.R. Barger, et al., Radiation plus Procarbazine, CCNU, and Vincristine in Low-Grade Glioma. N Engl J Med, 2016. 374(14): p. 1344–55.

8. Gerstl, J.V.E., M. Price, J.D. Bernstock, C. Kruchko, L. Spanehl, P.V. Karandikar, et al., Years of life lost due to central nervous system tumor subtypes in the United States. Neuro Oncol, 2025. 27(10): p. 2738–2746.

9. Ostrom, Q.T., D.J. Cote, M. Ascha, C. Kruchko, and J.S. Barnholtz-Sloan, Adult Glioma Incidence and Survival by Race or Ethnicity in the United States From 2000 to 2014. JAMA Oncol, 2018. 4(9): p. 1254–1262.

10. Wanis, H.A., H. Moller, K. Ashkan, and E.A. Davies, The Influence of Ethnicity on Survival from Malignant Primary Brain Tumours in England: A Population-Based Cohort Study. Cancers (Basel), 2023. 15(5).

11. Gabriel, A., J. Batey, J. Capogreco, D. Kimball, A. Walters, R.S. Tubbs, and M. Loukas, Adult brain cancer in the U.S. black population: a Surveillance, Epidemiology, and End Results (SEER) analysis of incidence, survival, and trends. Med Sci Monit, 2014. 20: p. 1510–7.

12. Patel, N.P., K.A. Lyon, and J.H. Huang, The effect of race on the prognosis of the glioblastoma patient: a brief review. Neurol Res, 2019. 41(11): p. 967–971.

13. Barnholtz-Sloan, J.S., J.L. Maldonado, V.L. Williams, W.T. Curry, E.A. Rodkey, F.G. Barker, 2nd, and A.E. Sloan, *Racial/ethnic differences in survival among elderly patients with a primary glioblastoma*. J Neurooncol, 2007. 85(2): p. 171–80.

14. Barnholtz-Sloan, J.S., A.E. Sloan, and A.G. Schwartz, Racial differences in survival after diagnosis with primary malignant brain tumor. Cancer, 2003. 98(3): p. 603–9.

15. Grossman, S.A., X. Ye, S. Piantadosi, S. Desideri, L.B. Nabors, M. Rosenfeld, et al., Survival of patients with newly diagnosed glioblastoma treated with radiation and temozolomide in research studies in the United States. Clin Cancer Res, 2010. 16(8): p. 2443–9.

16. Brat, D.J., K. Aldape, H. Colman, D. Figrarella-Branger, G.N. Fuller, C. Giannini, et al., cIMPACT-NOW update 5: recommended grading criteria and terminologies for IDH-mutant astrocytomas. Acta Neuropathol, 2020. 139(3): p. 603–608.

17. Ceccarelli, M., F.P. Barthel, T.M. Malta, T.S. Sabedot, S.R. Salama, B.A. Murray, et al., Molecular Profiling Reveals Biologically Discrete Subsets and Pathways of Progression in Diffuse Glioma. Cell, 2016. 164(3): p. 550–63.

18. Cancer Genome Atlas Research, N., D.J. Brat, R.G. Verhaak, K.D. Aldape, W.K. Yung, S.R. Salama, et al., Comprehensive, Integrative Genomic Analysis of Diffuse Lower-Grade Gliomas. N Engl J Med, 2015. 372(26): p. 2481–98.

19. Amirian, E.S., G.N. Armstrong, R. Zhou, C.C. Lau, E.B. Claus, J.S. Barnholtz-Sloan, et al., The Glioma International Case-Control Study: A Report From the Genetic Epidemiology of Glioma International Consortium. Am J Epidemiol, 2016. 183(2): p. 85–91.

20. Lim, H., M.C. Gingras, J. Zhao, J. Byun, P.D. Castro, S. Tsavachidis, et al., Somatic mutations of esophageal adenocarcinoma: a comparison between Black and White patients. Sci Rep, 2024. 14(1): p. 8988.

21. Rokita, J.L., K.S. Rathi, M.F. Cardenas, K.A. Upton, J. Jayaseelan, K.L. Cross, et al., Genomic Profiling of Childhood Tumor Patient-Derived Xenograft Models to Enable Rational Clinical Trial Design. Cell Rep, 2019. 29(6): p. 1675–1689 e9.

22. Tarasov, A., A.J. Vilella, E. Cuppen, I.J. Nijman, and P. Prins, Sambamba: fast processing of NGS alignment formats. Bioinformatics, 2015. 31(12): p. 2032–4.

23. McKenna, A., M. Hanna, E. Banks, A. Sivachenko, K. Cibulskis, A. Kernytsky, et al., The Genome Analysis Toolkit: a MapReduce framework for analyzing next-generation DNA sequencing data. Genome Res, 2010. 20(9): p. 1297–303.

24. Fan, Y., L. Xi, D.S. Hughes, J. Zhang, J. Zhang, P.A. Futreal, et al., MuSE: accounting for tumor heterogeneity using a sample-specific error model improves sensitivity and specificity in mutation calling from sequencing data. Genome Biol, 2016. 17(1): p. 178.

25. Manichaikul, A., J.C. Mychaleckyj, S.S. Rich, K. Daly, M. Sale, and W.M. Chen, Robust relationship inference in genome-wide association studies. Bioinformatics, 2010. 26(22): p. 2867–73.

26. Favero, F., T. Joshi, A.M. Marquard, N.J. Birkbak, M. Krzystanek, Q. Li, et al., Sequenza: allele-specific copy number and mutation profiles from tumor sequencing data. Ann Oncol, 2015. 26(1): p. 64–70.

27. Manders, F., A.M. Brandsma, J. de Kanter, M. Verheul, R. Oka, M.J. van Roosmalen, et al., MutationalPatterns: the one stop shop for the analysis of mutational processes. BMC Genomics, 2022. 23(1): p. 134.

28. Ostrom, Q.T., L. Bauchet, F.G. Davis, I. Deltour, J.L. Fisher, C.E. Langer, et al., The epidemiology of glioma in adults: a “state of the science” review. Neuro Oncol, 2014. 16(7): p. 896–913.

29. Wang, G.M., G. Cioffi, N. Patil, K.A. Waite, R. Lanese, Q.T. Ostrom, et al., Importance of the intersection of age and sex to understand variation in incidence and survival for primary malignant gliomas. Neuro Oncol, 2022. 24(2): p. 302–310.

30. Ostrom, Q.T., J.B. Rubin, J.D. Lathia, M.E. Berens, and J.S. Barnholtz-Sloan, Females have the survival advantage in glioblastoma. Neuro Oncol, 2018. 20(4): p. 576–577.

31. Zhao, Z., K.N. Zhang, Q. Wang, G. Li, F. Zeng, Y. Zhang, et al., Chinese Glioma Genome Atlas (CGGA): A Comprehensive Resource with Functional Genomic Data from Chinese Glioma Patients. Genomics Proteomics Bioinformatics, 2021. 19(1): p. 1–12.

32. Cote, D.J., Q.T. Ostrom, H. Gittleman, K.R. Duncan, T.S. CreveCoeur, C. Kruchko, et al., Glioma incidence and survival variations by county-level socioeconomic measures. Cancer, 2019. 125(19): p. 3390–3400.

33. Shabihkhani, M., D. Telesca, M. Movassaghi, Y.B. Naeini, K.M. Naeini, S.A. Hojat, et al., Incidence, survival, pathology, and genetics of adult Latino Americans with glioblastoma. J Neurooncol, 2017. 132(2): p. 351–358.

34. Shin, J.Y., J.K. Yoon, and A.Z. Diaz, Racial disparities in anaplastic oligodendroglioma: An analysis on 1643 patients. J Clin Neurosci, 2017. 37: p. 34–39.

35. Ostrom, Q.T., K.M. Egan, L.B. Nabors, T. Gerke, R.C. Thompson, J.J. Olson, et al., Glioma risk associated with extent of estimated European genetic ancestry in African Americans and Hispanics. Int J Cancer, 2020. 146(3): p. 739–748.

36. Yuan, J., Z. Hu, B.A. Mahal, S.D. Zhao, K.H. Kensler, J. Pi, et al., Integrated Analysis of Genetic Ancestry and Genomic Alterations across Cancers. Cancer Cell, 2018. 34(4): p. 549–560 e9.

37. Alexandrov, L.B., J. Kim, N.J. Haradhvala, M.N. Huang, A.W. Tian Ng, Y. Wu, et al., The repertoire of mutational signatures in human cancer. Nature, 2020. 578(7793): p. 94–101.

38. Zhang, B., W. Wan, Z. Li, Z. Gao, N. Ji, J. Xie, et al., A prognostic risk model for glioma patients by systematic evaluation of genomic variations. iScience, 2022. 25(12): p. 105681.

39. Daniel, P., S. Sabri, A. Chaddad, B. Meehan, B. Jean-Claude, J. Rak, and B.S. Abdulkarim, Temozolomide Induced Hypermutation in Glioma: Evolutionary Mechanisms and Therapeutic Opportunities. Front Oncol, 2019. 9: p. 41.

40. Li, C.H., S. Haider, and P.C. Boutros, Age influences on the molecular presentation of tumours. Nat Commun, 2022. 13(1): p. 208.

41. Weijers, D.D., S. Hinic, E. Kroeze, M.A. Gorris, G. Schreibelt, S. Middelkamp, et al., Unraveling mutagenic processes influencing the tumor mutational patterns of individuals with constitutional mismatch repair deficiency. Nat Commun, 2025. 16(1): p. 4459.

