## Supplemental for "Multi-Omics Study of Ancestry in Adults with Intracranial Cancers – Glioma (MOSAIC)"

**Manuscript title:**

**Supplementary Figures and Tables**

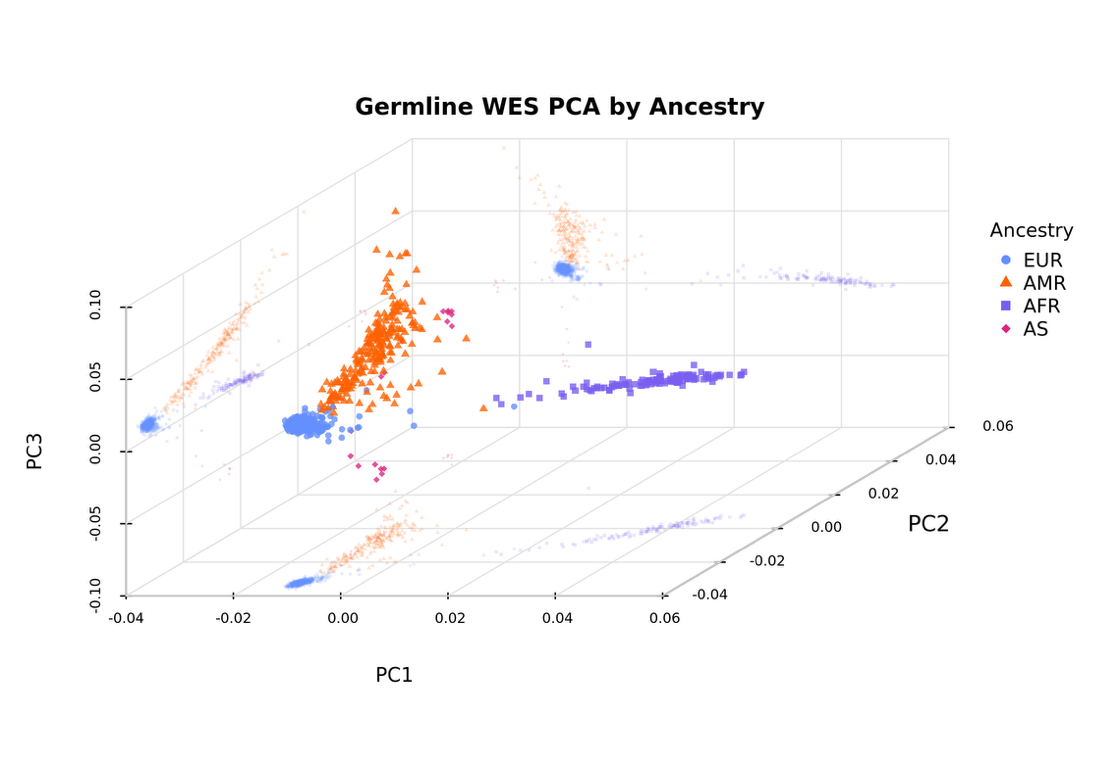

**Supplementary Figure 1. Germline WES PCA colored by genetic ancestry.**
Principal component analysis (PCA) of germline whole-exome sequencing (WES) data showing population structure across samples. Each point represents an individual sample projected onto the first three principal components (PC1, PC2, PC3). Samples are colored by inferred ancestry: European (EUR, blue), Admixed American (AMR, orange), African (AFR, purple), and Asian (AS, pink). Distinct clustering patterns are observed across ancestry groups, with clear separation along the principal component axes, reflecting underlying genetic variation and population stratification.

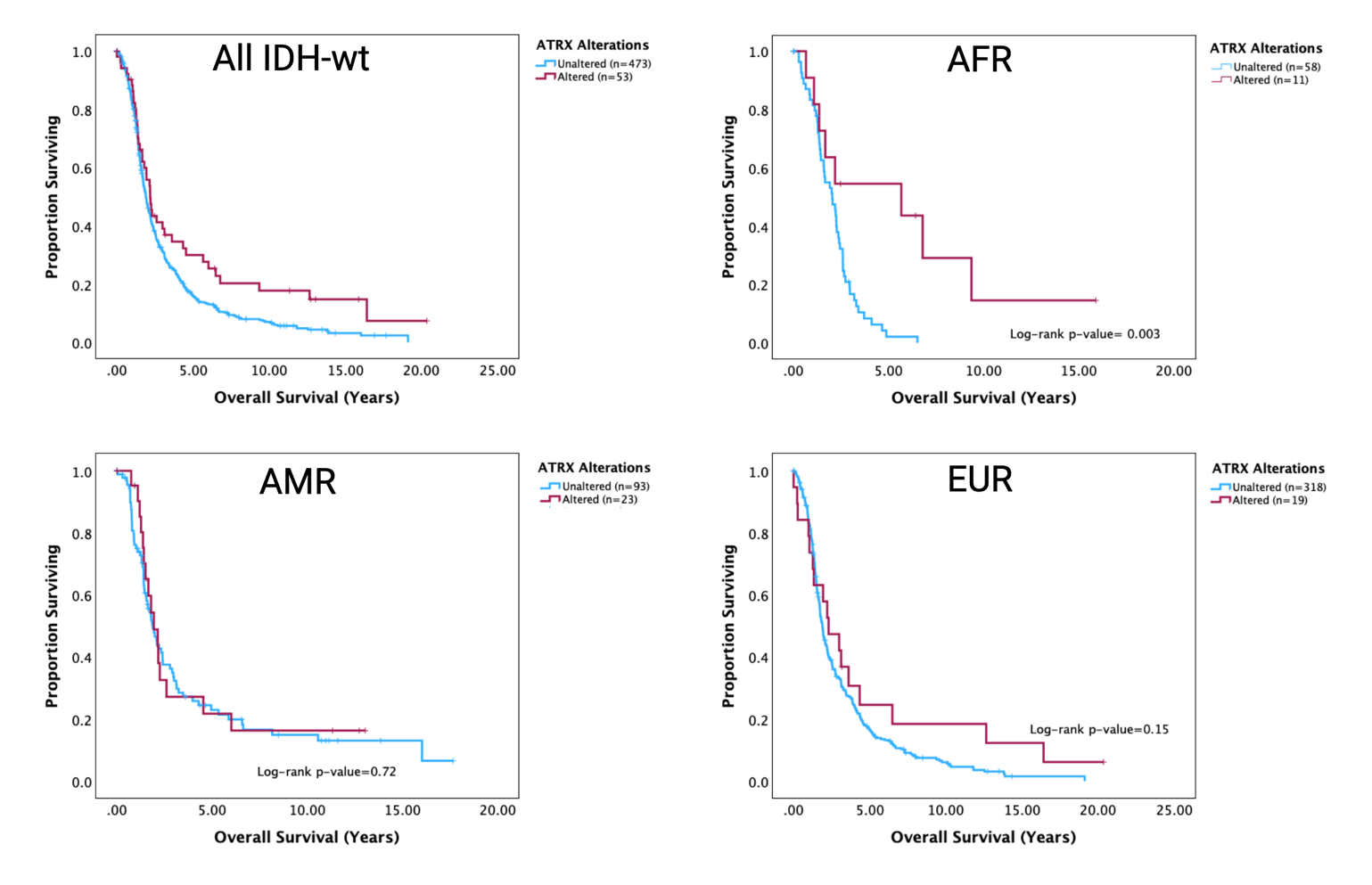

**Supplementary Figure 2. ATRX alterations are associated with improved survival in AFR but not AMR or EUR IDH-wildtype gliomas.**
Kaplan–Meier overall survival curves for patients with IDH-wildtype gliomas stratified by ATRX alteration status across ancestry groups: African (AFR), Admixed American (AMR), and European (EUR). Patients are grouped as ATRX-altered (red) and ATRX-unaltered (blue), with sample sizes indicated in each panel. In the AFR subgroup, ATRX alterations are associated with significantly improved overall survival (log-rank p = 0.003). No significant survival differences are observed in the AMR (p = 0.72) or EUR (p = 0.15) subgroups. These results suggest ancestry-specific prognostic relevance of ATRX alterations in IDH-wildtype glioma.

**Supplementary Table 1.** Signatures significantly associated with tumor type (Kruskal-Wallis test, FDR < 0.05).

| **Signature** | **P-value** | **Adjusted P-value** | **Etiology** | **Enriched in glioma subtype** |
| --- | --- | --- | --- | --- |
| SBS1 | 9.83e-56 | 5.90e-54 | Spontaneous deamination of 5-methylcytosine (clock-like) | IDH-wt |
| SBS87 | 4.41e-23 | 1.32e-21 | Thiopurine chemotherapy | Astro |
| SBS10b | 1.95e-08 | 3.89e-07 | POLE exonuclease domain mutations | Astro |
| SBS40 | 1.41e-07 | 2.11e-06 | Unknown (clock-like) | IDH-wt |
| SBS38 | 2.87e-06 | 3.44e-05 | Indirect effect of ultraviolet light | IDH-wt |
| SBS23 | 2.30e-05 | 2.30e-04 | Unknown | Astro |
| SBS37 | 4.28e-05 | 3.48e-04 | Unknown | Oligo |
| SBS19 | 4.65e-05 | 3.48e-04 | Unknown | Astro |
| SBS86 | 1.50e-04 | 9.97e-04 | Unknown chemotherapy | Astro |
| SBS4 | 2.12e-04 | 1.27e-03 | Tobacco smoking | IDH-wt |
| SBS32 | 5.47e-04 | 2.98e-03 | Azathioprine treatment | Astro |
| SBS6 | 6.42e-04 | 3.21e-03 | Defective DNA mismatch repair (MSI) | Astro |
| SBS44 | 1.97e-03 | 9.10e-03 | Defective DNA mismatch repair | IDH-wt |
| SBS3 | 2.55e-03 | 1.09e-02 | Defective homologous recombination DNA repair (BRCA1/2) | IDH-wt |
| SBS8 | 4.51e-03 | 1.81e-02 | Unknown (possible late replication errors) | IDH-wt |
| SBS11 | 8.59e-03 | 3.22e-02 | Temozolomide treatment | Oligo |
| SBS94 | 1.18e-02 | 4.15e-02 | Unknown | Astro |
| SBS7b | 1.31e-02 | 4.37e-02 | Ultraviolet light exposure | Oligo |

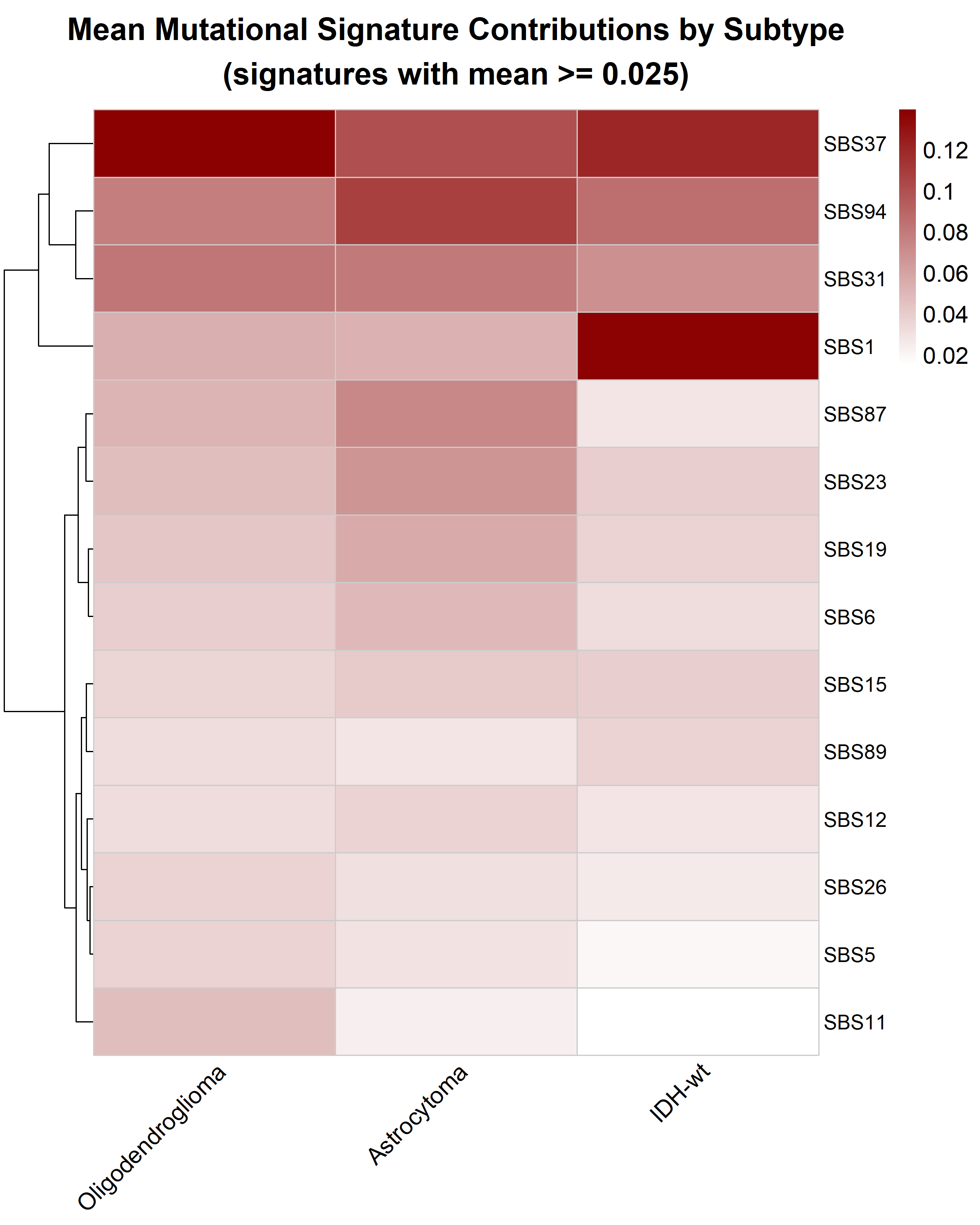

**Supplementary Figure 3.** **Heatmap of mean mutational signature contributions by tumor type.** Heatmap showing the mean contributions of COSMIC SBS v3.2 mutational signatures across glioma tumor types. Only signatures with a mean contribution ≥ 0.025 across tumor types are displayed to emphasize biologically relevant signals. Rows represent mutational signatures and columns represent tumor types, with color intensity indicating the average signature contribution. Hierarchical clustering of signatures highlights shared patterns of mutational processes across glioma subgroups.

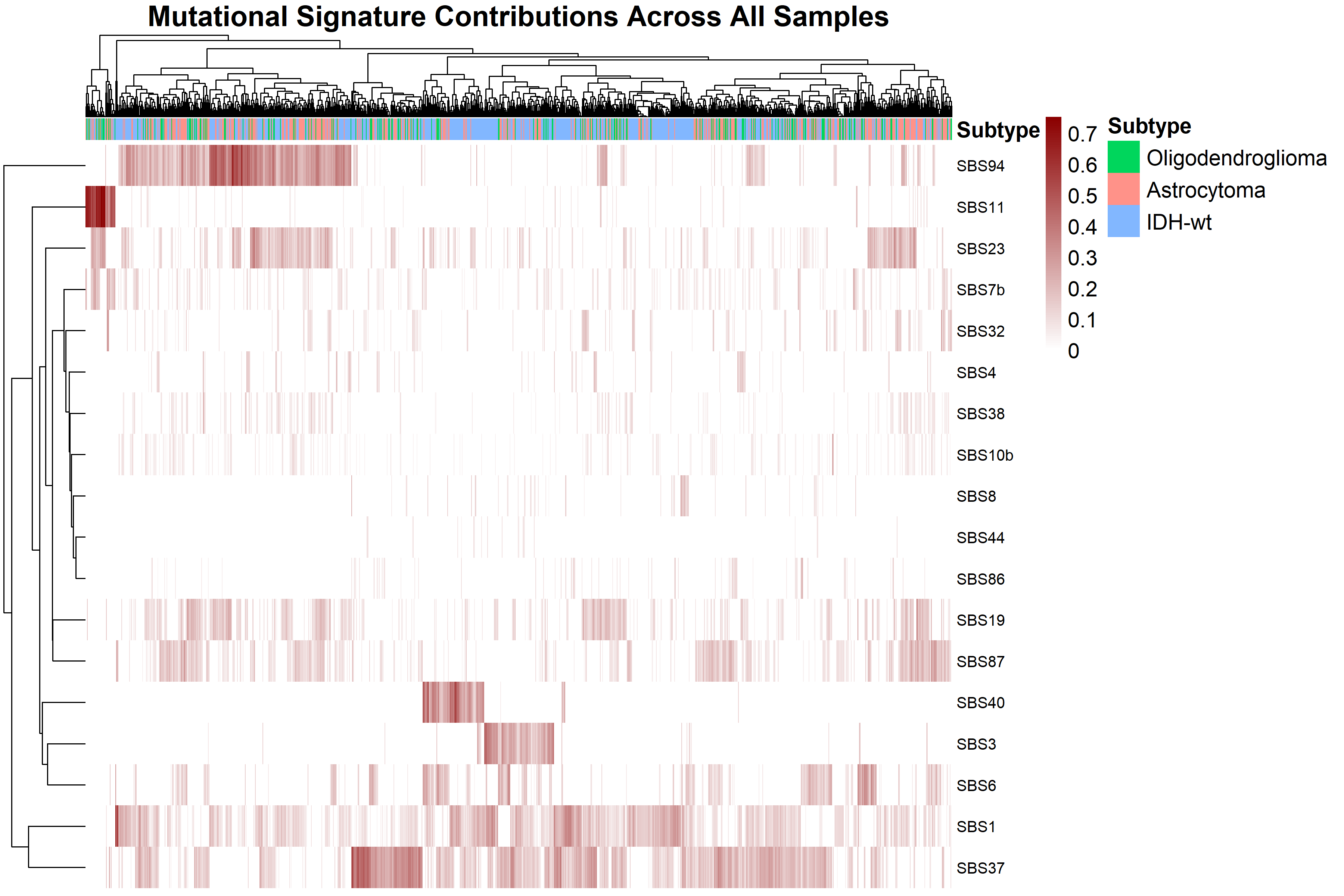

**Supplementary Figure 4.** **Mutational signature contributions across all glioma samples.**
Heatmap displaying contributions of COSMIC SBS mutational signatures across individual glioma samples. Rows represent mutational signatures and columns represent samples, ordered by tumor type as indicated in the top annotation. Color intensity reflects the relative contribution of each signature within a sample. Hierarchical clustering of signatures highlights patterns of co-occurrence and heterogeneity of mutational processes across the cohort.
